# Proton-mediated gating mechanism of anion channelrhodopsin-1

**DOI:** 10.1101/2021.07.22.453395

**Authors:** Masaki Tsujimura, Keiichi Kojima, Shiho Kawanishi, Yuki Sudo, Hiroshi Ishikita

**Affiliations:** Department of Applied Chemistry, The University of Tokyo, 7-3-1 Hongo, Bunkyo-ku, Tokyo 113-8654, Japan; Graduate School of Medicine, Dentistry and Pharmaceutical Sciences, Okayama University, Okayama 700-8530, Japan; Research Center for Advanced Science and Technology, The University of Tokyo, 4-6-1 Komaba, Meguro-ku, Tokyo 153-8904, Japan

**Author notes:** CORRESPONDING AUTHOR: Ishikita, Graduate School of Engineering, The University of Tokyo, 4-6-1 Komaba, Meguro-ku, Tokyo 153-8904, Japan, **E-mail:**.

## Abstract

Anion channelrhodopsin from *Guillardia theta* (*Gt*ACR1) has Asp234 (3.2 Å) and Glu68 (5.3 Å) near the protonated Schiff base. Here we investigate mutant *Gt*ACR1s (e.g., E68Q/D234N) expressed in HEK293 cells. The influence of the acidic residues on the absorption wavelengths were also analyzed, using a quantum mechanical/molecular mechanical approach. The calculated protonation pattern indicates that Asp234 is deprotonated and Glu68 is protonated in the original crystal structures. The D234E mutation and the E68Q/D234N mutation shortens and lengthens the measured and calculated absorption wavelengths, respectively, which suggests that Asp234 is deprotonated in the wild type *Gt*ACR1. Molecular dynamics simulations show that upon mutation of deprotonated Asp234 to asparagine, deprotonated Glu68 reorients towards the Schiff base and the calculated absorption wavelength remains unchanged. The formation of the proton transfer pathway via Asp234 toward Glu68 and the disconnection of the anion conducting channel are likely a basis of the gating mechanism.

## INTRODUCTION

Anion channelrhodopsins (ACRs) are light-gated anion channels that undergo photoisomerization at the retinal chromophore, which is covalently attached to a conserved lysine residue via the protonated Schiff base, from all-*trans* to 13-*cis*. Natural ACRs were identified in the cryptophyte *Guillardia theta* (*Gt*ACR1 and *Gt*ACR2) ^1^. ACRs hyperpolarize the membrane through anion-import and can widely be used as neural silencing tools in optogenetics ^2,3^. Microbial rhodopsins have acidic residues or Cl^−^ at the Schiff base moiety to stabilize the protonated Schiff base as counterions. Counterions play a major role in determining the absorption wavelength and the function of the protein ^4^. The X-ray crystal structures of ACR from *Guillardia theta* (*Gt*ACR1) show that two acidic residues, Glu68 and Asp234, exist at the corresponding positions (Figure 1) ^5,6^.

**Figure 1.**
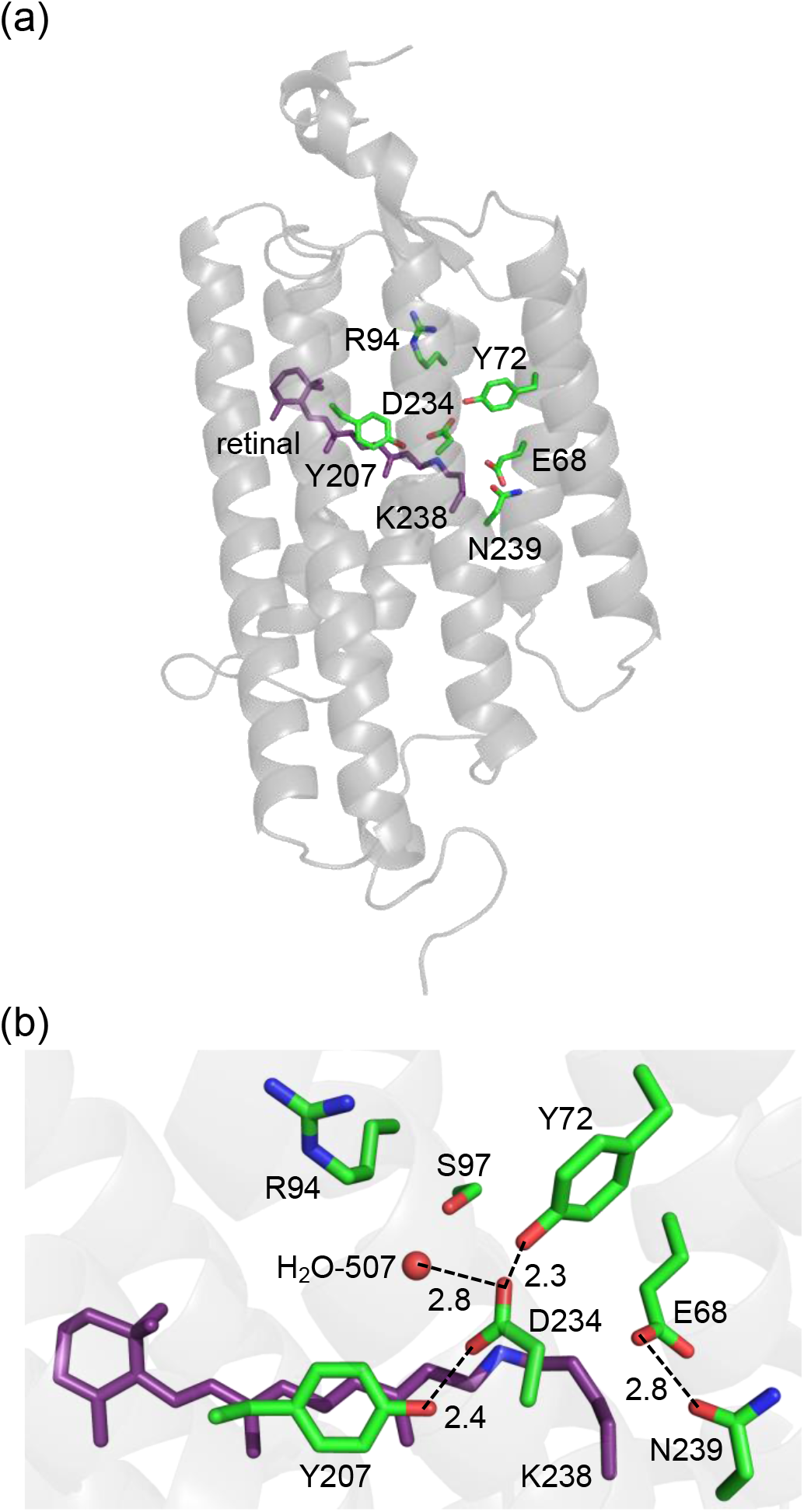
(a) Overview of the *Gt*ACR1 structure. (b) Residues near the Schiff base binding site in *Gt*ACR1 (PDB code: 6EDQ ^6^).

It was proposed that both Glu68 and Asp234 were protonated in *Gt*ACR1 ^5,7–9^ in contrast to other microbial rhodopsins, because the absorption wavelengths remain unchanged upon the E68Q and D234N mutations ^5,7,8^. Indeed, the C=C stretching frequency of the retinal is not significantly affected upon the E68Q and D234N mutations in resonance Raman spectroscopy, which implies that the electrostatic interaction between the retinal and protein environment remains unchanged ^8^. In addition, the C=O stretching frequency for a protonated carboxylate, which is observed in the wild type *Gt*ACR1, disappears in the E68Q ^10,11^ and D234N ^5^ *Gt*ACR1s according to Fourier-transform infrared (FTIR) spectroscopy analysis.

Nevertheless, it is an open question whether Asp234 is protonated. Kim et al. pointed out that the loss of photocurrent in the D234N *Gt*ACR1 cannot be easily understood if Asp234 is protonated, as the influence of the mutation of protonated aspartate to asparagine on the protein function is often small ^5^. *Gt*ACR1 crystal structures show that the protein electrostatic environment at the Schiff base moiety is highly conserved between *Gt*ACR1 and bacteriorhodopsin (BR). Tyr57, Arg82, Tyr185, and Lys216, which are responsible for the low p*K*_a_ of −2 for the counterion Asp212 in BR ^12^, are fully conserved as Tyr72, Arg94, Tyr207, and Lys238 in *Gt*ACR1. According to Li et al., Tyr72 and Tyr207 donate H-bonds to Asp234 in *Gt*ACR1 ^6^: this suggests that deprotonated Asp234 is stabilized, decreasing p*K*_a_(Asp234), as observed for deprotonated Asp212 in BR. In addition, resonance Raman spectroscopy analysis indicates that the Schiff base Lys238 is also protonated ^8^ as observed in other microbial rhodopsins. The presence of the positively charged Schiff base needs to have an adjacent negative charge (e.g., deprotonated acidic residue) to effectively decrease the energy in *Gt*ACR1. To the best of our knowledge, microbial rhodopsins have more than one deprotonated acidic residue adjacent to the retinal Schiff base (e.g., ^4^). This also holds true for channelrhodopsin from *Chlamydomonas noctigama* (Chrimson) and rhodopsin phosphodiesterase (Rh-PDE), which have both deprotonated and protonated acidic residues near the Schiff base ^13,14^. That is, either Glu68 or Asp234 may be deprotonated in *Gt*ACR1. Deprotonation of Asp234 is energetically more favorable than deprotonation of Glu68 in the presence of protonated Schiff base, as the electrostatic interaction with Asp234 (3.2 Å) is larger than with Glu68 (5.3 Å). Alternatively, Cl^−^ may exist and act as a counterion, as observed in Cl^−^ pumping rhodopsins ^15,16^. However, the corresponding electron density is not observed in *Gt*ACR1 ^5,6^. In addition, no spectral changes are reported upon deionization of the sample or exchange from Cl^−^ to SO_4_^2–^ buffer ^7^. So far, the counterion of *Gt*ACR1 remains unknown.

Recently, Dreier et al. proposed that Asp234 is deprotonated in the dark and act as a counterion according to FTIR measurements and molecular dynamics (MD) simulations^11^. The C=O stretching frequencies of 1740 (–)/1732 (+) cm^−1^ for protonated Asp234 at 77 K observed by Kim et al. ^5^, were not observed at 293 K by Dreier et al. ^11^. In addition, MD simulations indicated that the H-bond network that involves Tyr72, Tyr207, and Asp234 was stable with deprotonated Asp234 but unstable with protonated Asp234 ^11^. Indeed, the presence of deprotonated Asp234 was already suggested based on the homology modeling of *Gt*ACR2 ^17^ before the crystal structures of *Gt*ACR1 were reported.

*Gt*ACR1 undergoes a photocycle including K, L, M, N, and O intermediates (Figure 2) ^7^. The L-state represents the anion conducting state. The L- to M-state transition involves the deprotonation of the Schiff base and the photocurrent decay. The fast photocurrent decay (fast channel closing) corresponds to the M-state formation (i.e. proton release from the Schiff base), and the slow photocurrent decay corresponds to the M-state decay ^7,18^ (Figure 2). Glu68 is likely to accept a proton from the Schiff base upon the M-state formation, as a decrease in the accumulation of the M-state was observed in the E68Q *Gt*ACR1 ^7^. However, it remains unclear whether Glu68 is the initial proton acceptor in the wild type *Gt*ACR1. The *Gt*ACR1 crystal structures show that Asp234 is closer to the Schiff base (Figure 1) ^5,6,19^. In addition, the E68Q mutation did not completely inhibit the M-state formation, which indicates that alternative proton accepter exists ^7^.

**Figure 2.**
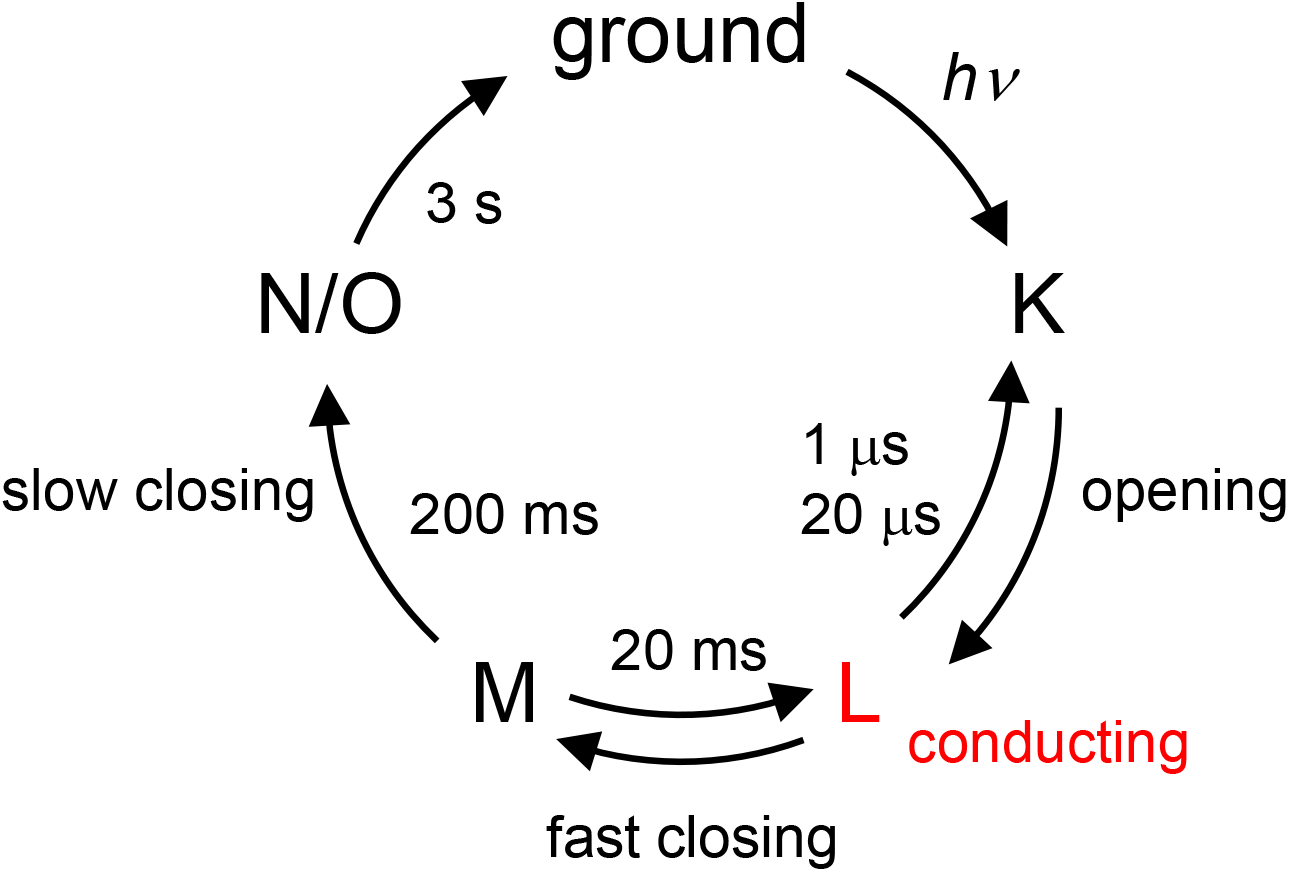
Photocycle of *Gt*ACR1 proposed in ref. ^7^. The L- to M-state and M- to N/O- state transitions involve deprotonation and reprotonation of the Schiff base, respectively.

To identify counterion and clarify the proton-mediated gating mechanism of *Gt*ACR1, we investigate the Glu68 and Asp234 mutant proteins (E68Q, E68D, D234N, D234E, and E68Q/D234N) expressed in HEK293 cells. The protonation states are calculated by solving the Poisson-Boltzmann equation and evaluated by conducting MD simulations. Using a quantum mechanical/molecular mechanical (QM/MM) approach, the absorption wavelengths are calculated and the microscopic origin of the wavelength shifts upon the mutations are analyzed.

## RESULTS

### Protonation states of Glu68 and Asp234

The calculated protonation pattern (Table 1) and p*K*_a_ values (Tables 2 and 3) show that Asp234 is deprotonated whereas Glu68 is protonated in the *Gt*ACR1 crystal structures ^5,6^ (Table 1 and Table S1). The calculated protonation pattern shows that Asp234 is deprotonated in the wild type *Gt*ACR1 even using the MD-generated conformations with protonated Asp234 (Table 1). These results suggest that deprotonation of Asp234, “the only residue directly interacting with the protonated Schiff base ^6^”, is a prerequisite to stabilize the protonated Schiff base, as suggested by Dreier et al. ^11^.

**Table 1.** Protonation states of Glu68 and Asp234. Changes in the protonation states are in bold. –, not applicable.

|  |  | Fixed state<br>during MD | Calculated state<br>after MD <sup>a</sup> |
| --- | --- | --- | --- |
| wild type (1) | E68 | protonated | protonated |
|  | D234 | deprotonated | deprotonated |
| wild type (2) | E68 | protonated | protonated |
|  | D234 | protonated | **deprotonated** |
| E68D (1) | D68 | protonated | protonated |
|  | D234 | deprotonated | deprotonated |
| E68D (2) | D68 | deprotonated | **protonated** |
|  | D234 | deprotonated | deprotonated |
| E68Q | Q68 | – | – |
|  | D234 | deprotonated | deprotonated |
| D234E | E68 | protonated | protonated |
|  | E234 | deprotonated | deprotonated |
| D234N | E68 | deprotonated | deprotonated |
|  | N234 | – | – |
| E68Q/D234N | Q68 | – | – |
|  | N234 | – | – |

<sup>a</sup> Protonation patterns obtained using the MD-generated conformations.

**Table 2.**
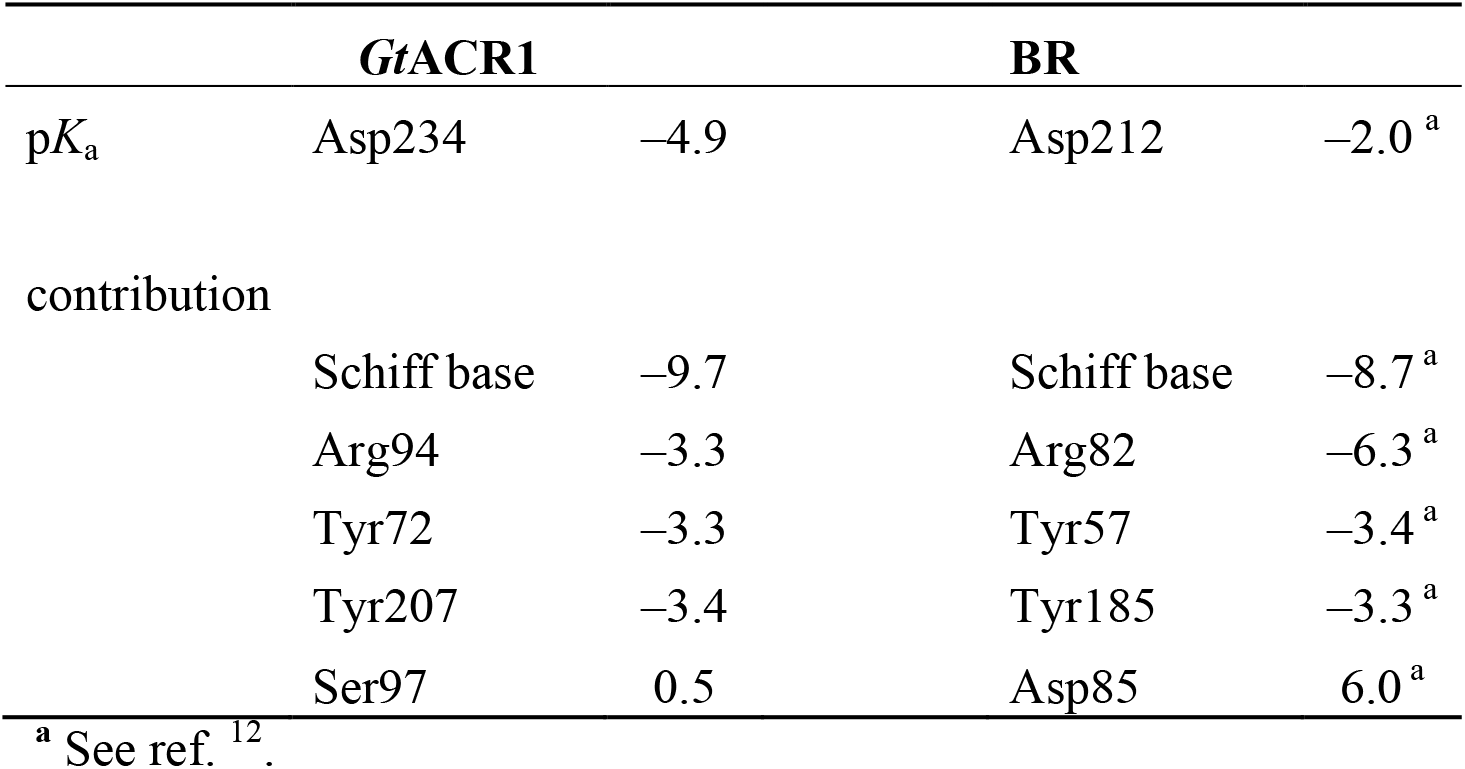
p*K*_a_ for Asp234 in *Gt*ACR1 and Asp212 in BR.

|  | ***GtACR1*** |  | **BR** |  |
| --- | --- | --- | --- | --- |
| $pK_a$ | Asp234 | −4.9 | Asp212 | −2.0<sup>a</sup> |
| contribution |  |  |  |  |
|  | Schiff base | −9.7 | Schiff base | −8.7<sup>a</sup> |
|  | Arg94 | −3.3 | Arg82 | −6.3<sup>a</sup> |
|  | Tyr72 | −3.3 | Tyr57 | −3.4<sup>a</sup> |
|  | Tyr207 | −3.4 | Tyr185 | −3.3<sup>a</sup> |
|  | Ser97 | 0.5 | Asp85 | 6.0<sup>a</sup> |

<sup>a</sup> See ref.<sup>12</sup>.

**Table 3.** p*K*_a_ for Glu68 in *Gt*ACR1. The corresponding sites of BR are shown in the same line. –, not applicable.

|  | *Gt*ACR1 |  | BR |  |
| --- | --- | --- | --- | --- |
| p*K*<sub>a</sub> | Glu68 | 12 | Ala53 | – |
| contribution |  |  |  |  |
|  | Schiff base | –6.6 | Schiff base | – |
|  | Arg94 | –1.3 | Arg82 | – |
|  | Asn239 | 2.5 | Val217 | – |
|  | Asp234 | 4.9 | Asp212 | – |

<sup>a</sup> See ref. <sup>12</sup>.

p*K*_a_(Asp234) = −5 (Table 2) is significantly low and even lower than p*K*_a_(Asp212) = −2 in BR ^12^. The crystal structures show that the protein electrostatic environment at the Schiff base moiety is highly conserved between *Gt*ACR1 and BR. Tyr72 and Tyr207 donate H-bonds to each carboxyl O site of Asp234 in *Gt*ACR1 ^6,11^, whereas Tyr57 and Tyr185 donate H-bonds to each carboxyl O site of Asp212 in BR ^12^. Thus, each tyrosine residue stabilizes the deprotonated state of Asp234, decreasing p*K*_a_(Asp234) in *Gt*ACR1 by ∼3 (Table 2), as observed in BR ^12^. The tendency is also observed for the conserved residue pairs, Arg94/Arg82 and Lys238/Lys216 in *Gt*ACR1/BR (Table 2). Asp85, which increases p*K*_a_(Asp212) in BR by 6, is replaced with Ser97, which has no influence on p*K*_a_(Asp234) in *Gt*ACR1 (Table 2). This discrepancy contributes to the low p*K*_a_(Asp234) in *Gt*ACR1, which is lower than p*K*_a_(Asp212) in BR. As far as the original geometry of the *Gt*ACR1 crystal structure is analyzed, no residue that increases p*K*_a_(Asp234) significantly is identified (Table 2).

Recently, Li et al. reported the *Gt*ACR1 conformation (pre-activating state), where Arg94 forms a salt-bridge with Asp234 ^19^. The influence of Arg94 on p*K*_a_(Asp234) (∼3) indicates that the electrostatic link between Arg94 and Asp234 exists even in the ground state (Table 2). It seems possible that the electrostatic interaction between deprotonated Asp234 and channel-gating Arg94 (Table 2) is absent in the D234N *Gt*ACR1, leading to the loss of the photocurrent ^5^.

In contrast, p*K*_a_(Glu68) is high, 12 (Table 3), which is consistent with the reported protonation state of Glu68 ^8,10,11^. The high p*K*_a_(Glu68) value can be primarily due to the presence of anionic Asp234 whose deprotonated state is stabilized by the protonated

Schiff base (Table 3). Charge neutral Ala53 exists at the corresponding position in BR, which is also consistent with the protonation of Glu68 (Table 3).

Exceptionally, Glu68 is deprotonated only in the D234N *Gt*ACR1. The influence of the protonated Schiff base is weaker on Glu68 than on Asp234 (Tables 2 and 3), which allows to stabilize the putative protonated Glu68 conformation in MD simulations (Table 1). However, the experimentally measured absorption wavelength cannot be reproduced unless Glu68 is deprotonated in the D234N *Gt*ACR1 (Table 4). This is consistent with the absence of the 1708 cm^−1^ band in the D234N *Gt*ACR1, which is assigned to protonated Glu68 in FTIR measurements ^11^. These results confirm that the presence of a negative charge at the protonated Schiff base moiety is a prerequisite to stabilize the protonated Schiff base, as observed in other microbial rhodopsins ^4^, including Chrimson and Rh-PDE ^13,14^.

**Table 4.**
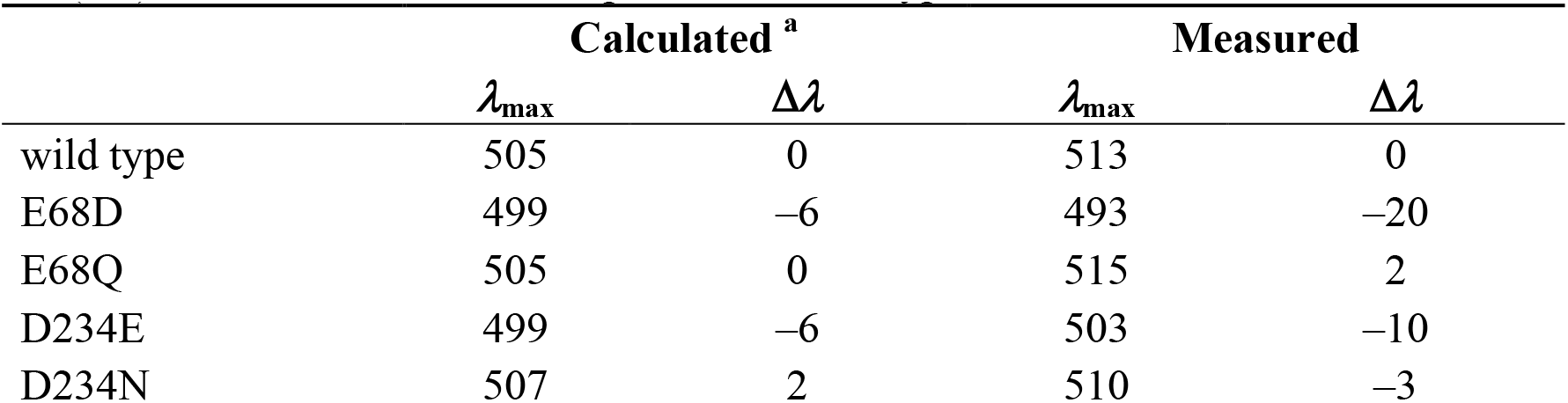
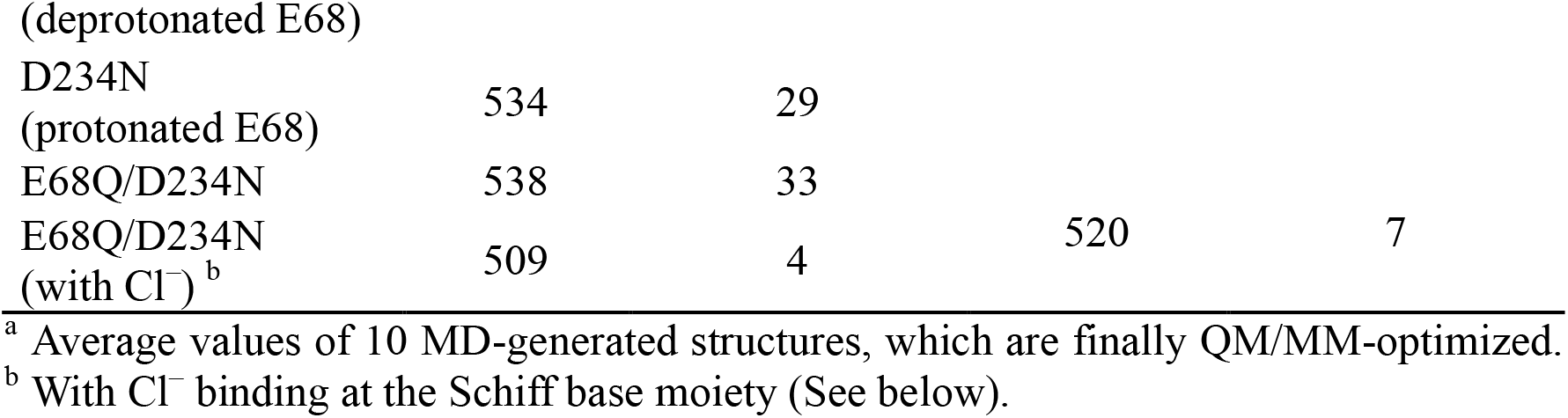
Calculated and experimentally measured absorption wavelengths *λ*_max_ (nm). Δ*λ* (nm) denotes the shift with respect to the wild type *Gt*ACR1.

### Absorption wavelengths of the Glu68 and Asp234 mutant *Gt*ACR1s

The experimentally measured absorption wavelengths for the Glu68 and Asp234 mutant proteins (E68Q, E68D, D234N, D234E, and E68Q/D234N) expressed in HEK293 cells are shown in Figure 3.

**Figure 3.**
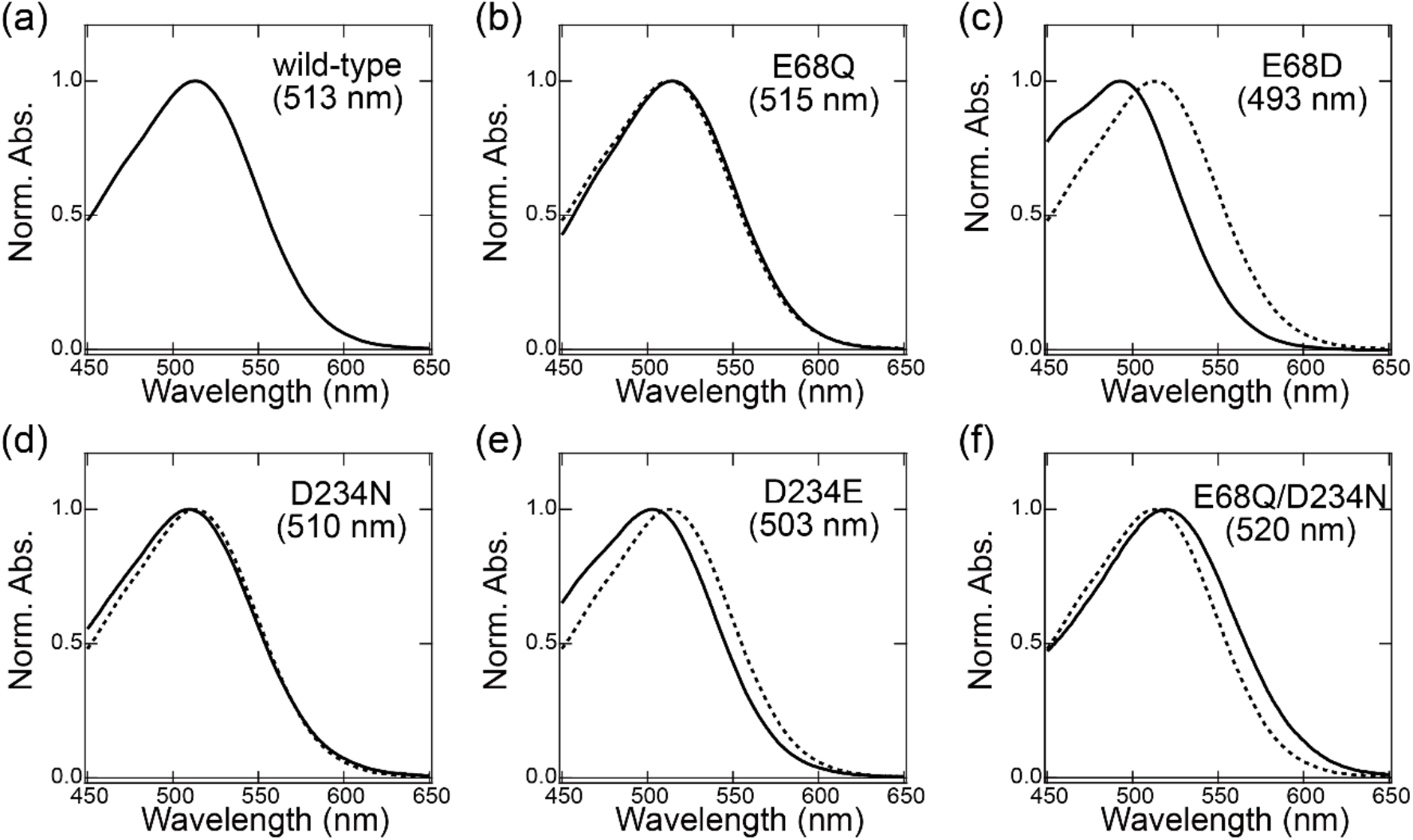
Absorption spectrum of (a) wild type, (b) E68Q, (c) E68D, (d) D234N, (e) D234E, and (f) E68Q/D234N *Gt*ACR1s in DDM micelles. Spectra are normalized at peak absorbance. The spectra for the wild type (dotted curves in panels b to f) are shown for comparison.

In the present study, the experimentally measured absorption wavelengths are the same for the wild type and D234N *Gt*ACR1s (Table 4), which is consistent with results reported previously ^5,7,8^. Notably, the calculated absorption wavelengths are also the same for the wild type and D234N *Gt*ACR1s irrespective of deprotonated Asp234 in the wild type *Gt*ACR1 (Table 4). This can be explained as, in the D234N *Gt*ACR1, deprotonated Glu68 moves toward the positively charged Schiff base, which fully substitutes a role of deprotonated Asp234 in stabilizing the positively charged Schiff base (Figure 4a, b, and Table 5). A similar conformation of Glu68, which orients toward the Schiff base, was previously reported for the corresponding residues of *Gt*ACR2 (Glu64) ^17^ and channelrhodopsin from *Chlamydomonas reinhardtii* (Glu90) ^20^. Thus, the absence of the change in the absorption wavelength upon D234N mutation ^5^ does not necessarily indicate that Asp234 is protonated in the wild type *Gt*ACR1.

**Figure 4.**
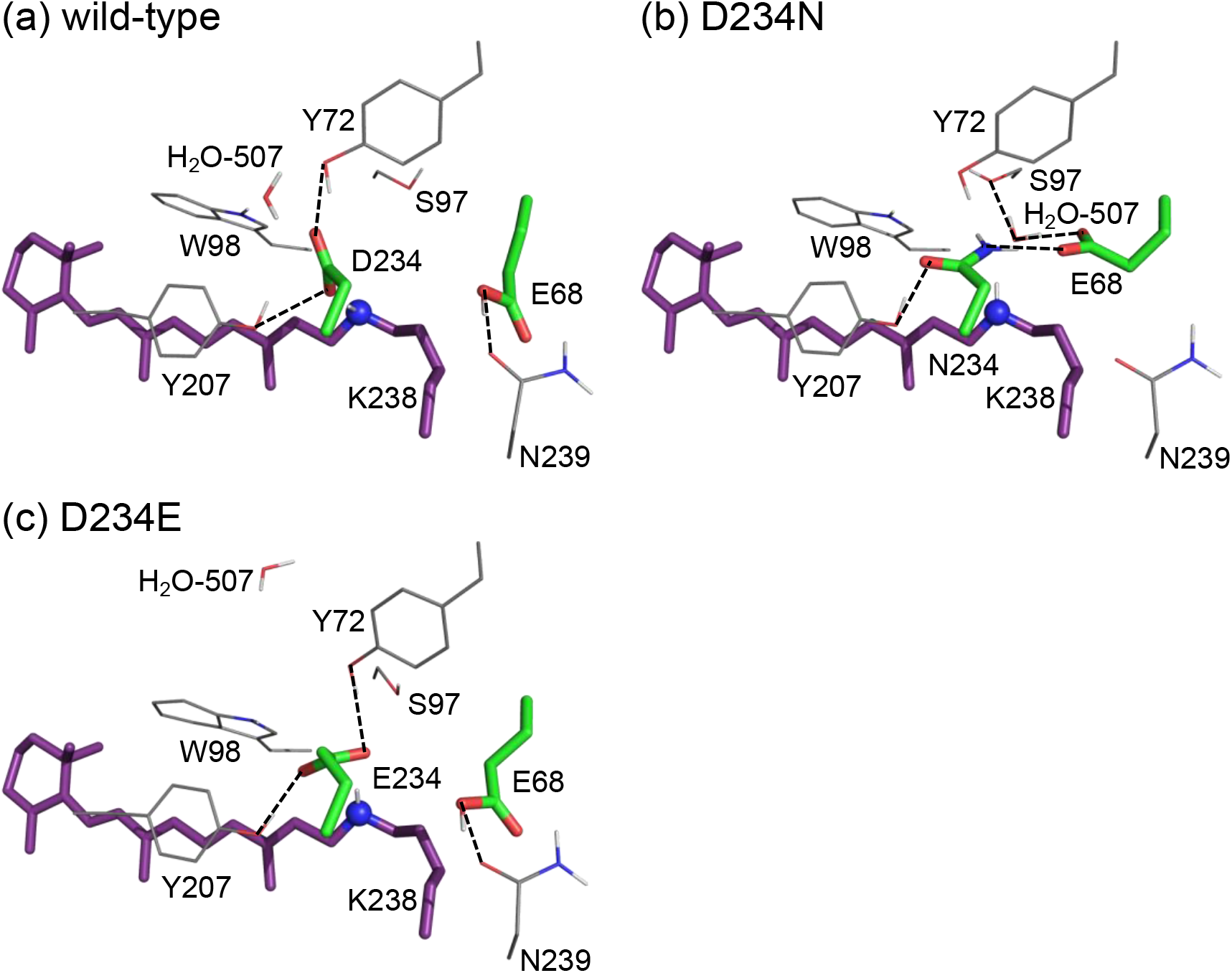
The H-bond network of the Schiff base of (a) wild type, (b) D234N, and (c) D234E *Gt*ACR1s. Representative MD-generated conformations, whose absorption wavelengths are closest to the average value for the 10 MD-generated structures, are shown. Dotted lines indicate H-bonds.

**Table 5.**
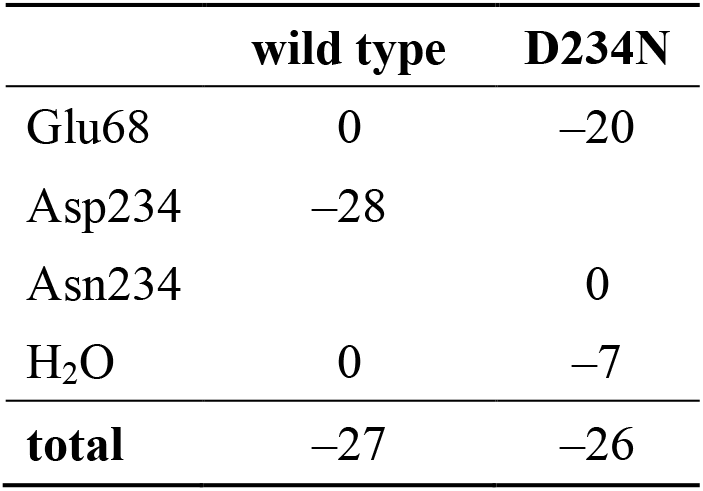
Components that contribute to the absorption wavelength in the wild type and D234N *Gt*ACR1s (nm). The contributions were analyzed using a single MD-generated structure whose absorption wavelength is closest to the average value of the 10 MD-generated structures.

|  | wild type | D234N |
| --- | --- | --- |
| Glu68 | 0 | -20 |
| Asp234 | -28 |  |
| Asn234 |  | 0 |
| H<sub>2</sub>O | 0 | -7 |
| **total** | **-27** | **-26** |

The existence of deprotonated Asp234 in the wild type *Gt*ACR1 can also be understood from the absorption wavelength in the D234E *Gt*ACR1. The distance between Glu234 and the Schiff base (2.9 Å in MD-generated conformations) in the D234E *Gt*ACR1 is shorter than that between Asp234 and the Schiff base (3.6 Å in MD-generated conformations) in the wild type GtACR1, because glutamate is longer than aspartate (Figure 4a, c). The absorption wavelength is short as the electrostatic interaction between the deprotonated counterion and the protonated Schiff base is strong ^4^. Remarkably, the D234E mutation leads to a decrease in the absorption wavelength (Table 4, Figure 3), which suggests that Asp234 is deprotonated in the wild type *Gt*ACR1. From the same analogy, the decrease in the absorption wavelength upon the E68D mutation (Figure 3, Table 4) is due to protonated Glu68 in the wild type *Gt*ACR1.

As far as we are aware, the absorption wavelength of the isolated E68Q/D234N *Gt*ACR1 is not reported (e.g., ^7,11^). We successfully isolated a photoactive form of E68Q/D234N *Gt*ACR1 using the HEK293 cell expression system, which has been widely used for the functional expression in animal rhodopsins ^21,22^. The experimentally measured absorption wavelength in the isolated E68Q/D234N protein is 7 nm longer than that in the wild type protein (Figure 3), which indicates that Glu68 or Asp234 must be deprotonated in the wild type *Gt*ACR1. As Asp234 is closer to the Schiff base (3.2 Å) than Glu68 (5.3 Å) ^6^, it seems more likely that Asp234 is deprotonated in the wild type *Gt*ACR1.

Microbial rhodopsins, including Chrimson and Rh-PDE ^13,14^, have more than one deprotonated acidic residue adjacent to the retinal Schiff base ^4^. The loss of two acidic residues upon the E68Q/D234N mutation requires an additional negative charge as far as the Schiff base remains protonated. Thus, it seems possible that Cl^−^ exist to stabilize the protonated Schiff base specifically in the E68Q/D234N *Gt*ACR1, because the next closest acidic residue, Glu60, is 10 Å away from the Schiff base. The presence of Cl^−^ in the E68Q/D234N *Gt*ACR1 is not reported. To investigate the existence of Cl^−^, isolated E68Q/D234N samples were solubilized in Cl^−^-free buffer. However, denaturation of the samples did not allow us to conclude the existence of Cl^−^. In QM/MM calculations and MD simulations, the binding of Cl^−^ at Thr71/Asn234 or Ser97/Lys238 is more stable in the E68Q/D234N *Gt*ACR1 than in the wild type protein (Figure S1, Table S2). The increase in the calculated absorption wavelength upon the E68Q/D234N mutation (33 nm) is overestimated in the absence of Cl^−^, whereas the corresponding increase (4 nm) is at the same level as that measured experimentally in the presence of Cl^−^ (Table 4). Thus, Cl^−^ is likely to exist near the protonated Schiff base to compensate for the loss of two acidic residues in the E68Q/D234N *Gt*ACR1.

The E68Q mutation does not alter the absorption wavelength (Table 4) as reported previously ^7,8^, thereby suggesting that Glu68 is protonated in the presence of deprotonated Asp234 (e.g., wild type *Gt*ACR1) (Table 1).

## DISCUSSION

Overall, the present results show that Asp234 is deprotonated in the wild type *Gt*ACR1, as indicated by the following findings: i) The E68Q/D234N mutation leads to an increase in the absorption wavelength (Table 4), which indicates that Glu68 or Asp234 is deprotonated in the wild type *Gt*ACR1 (Table 1); ii) The absorption wavelength in the D234E *Gt*ACR1 is shorter than in the wild type protein (Table 4), which can be explained only by the presence of a deprotonated acidic residue (Table 1); iii) Glu68, which is protonated in the wild type *Gt*ACR1, is deprotonated in the D234N *Gt*ACR1 (Table 1). If Glu68 remained protonated in the D234N *Gt*ACR1, the absorption wavelength would be significantly longer as compared with the wild type *Gt*ACR1 (Table 4). In any *Gt*ACR1, a negative charge needs to be present as far as the Schiff base is protonated; iv) Mutation of deprotonated Asp234 to uncharged asparagine does not alter the calculated absorption wavelength (Tables 4 and 5, Figure 4). Thus, the absence of changes in the absorption wavelength upon the D234N mutation does not serve as a basis of the presence of protonated Asp234 in the wild type *Gt*ACR1.

Based on these findings, loss of photocurrent upon the D234N mutation ^5^ can be due to loss of Asp234, which is deprotonated in the wild type *Gt*ACR1. It seems possible that the electrostatic interaction between deprotonated Asp234 and channel-gating Arg94 ^19^ (Table 2) is absent in the D234N *Gt*ACR1, leading to loss of the photocurrent ^5^. Intriguingly, MD simulations show that upon the D234N mutation, deprotonated Glu68 reorients toward and interferes with the channel bottle neck (Figure 5). It seems likely that Glu68 acts as a proton acceptor, forming the M-state (i.e., fast channel closing), in the D234N *Gt*ACR1 ^7^, as deprotonated Glu68 is sufficiently close to the protonated Schiff base (Figure 4b). The reorientation of deprotonated Glu68 toward the protonated Schiff base and the interference with the channel bottle neck may also explain why the photocurrent is abolished ^5^ irrespective of the accumulation of the M-state ^7^ (i.e., with deprotonated Schiff base) in the D234N *Gt*ACR1. Thus, not only the gating (Arg94) but also the conduction (Glu68) can be inhibited in the D234N *Gt*ACR1.

**Figure 5.**
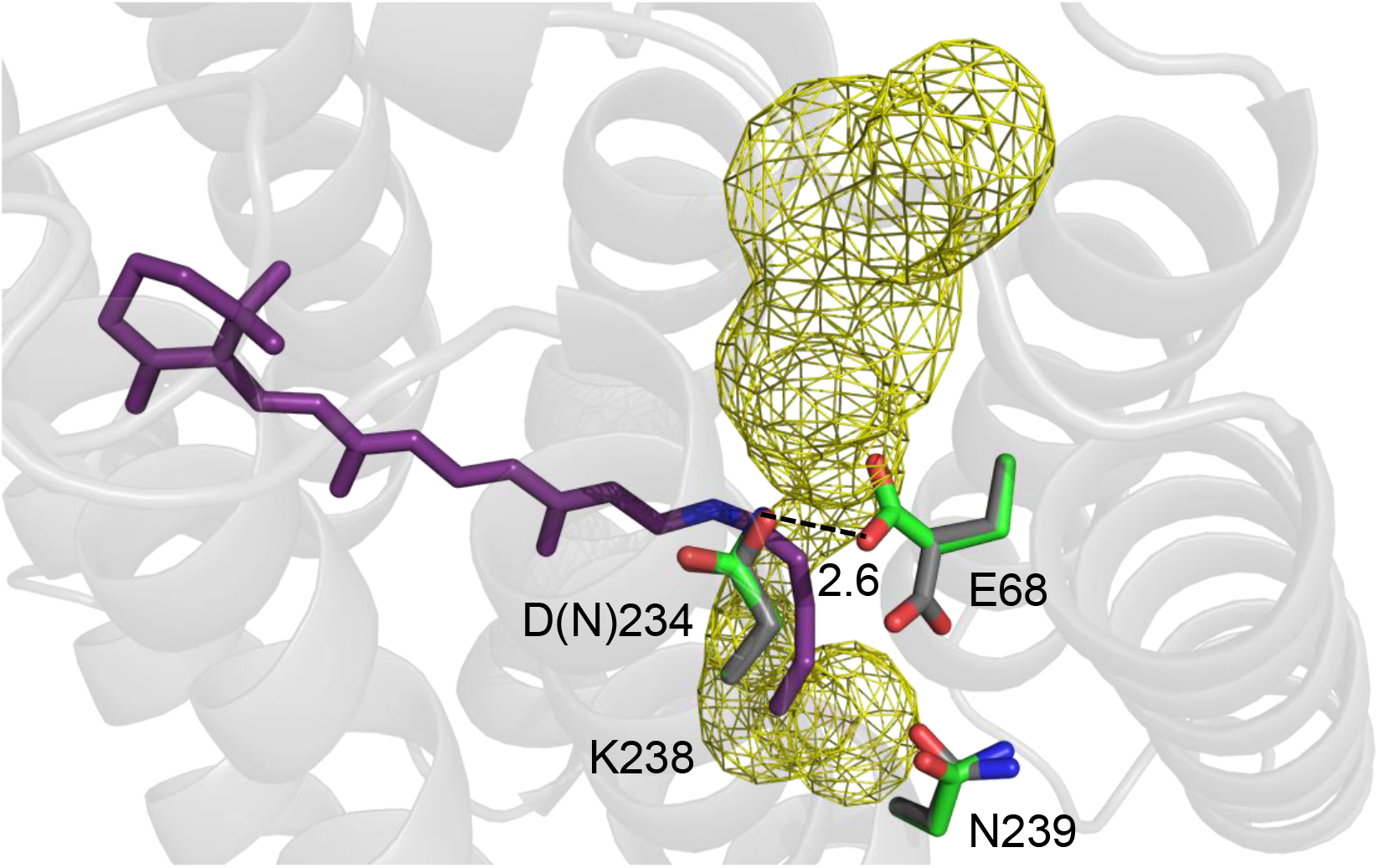
Channel space and side-chain orientations in the wild type (gray sticks, PDB code 6EDQ ^6^) and D234N (green sticks, a representative MD-generated conformation) *Gt*ACR1s. The yellow mesh indicates the channel space in the wild type *Gt*ACR1 analyzed using the CAVER program ^23^. Note that the channel space is consistent with that reported by Li et al. ^6^.

The formation of an H-bond between Asn234 and deprotonated Glu68 in the D234N *Gt*ACR1 (Figure 5) also suggests that Glu68 accepts the proton from protonated Asp234 in the M-state of the wild type *Gt*ACR1 (Figure 6). Then, the absence of Glu68 as a proton acceptor of Asp234 may affect the release of the proton from the Schiff base in the E68Q *Gt*ACR1. Indeed, it has been reported that the E68Q mutation affects the channel gating mechanism ^18^, leading to a decrease in the M-state (i.e., deprotonated Schiff base) accumulation ^7^. The conformation of Glu68 as a proton acceptor of Asp234 interferes with the channel bottle neck (Figure 5). This may explain why the M-state formation (i.e., release of the proton from the Schiff base via Asp234 to Glu68) corresponds to the fast channel closing (Figures 2 and 6).

**Figure 6.**
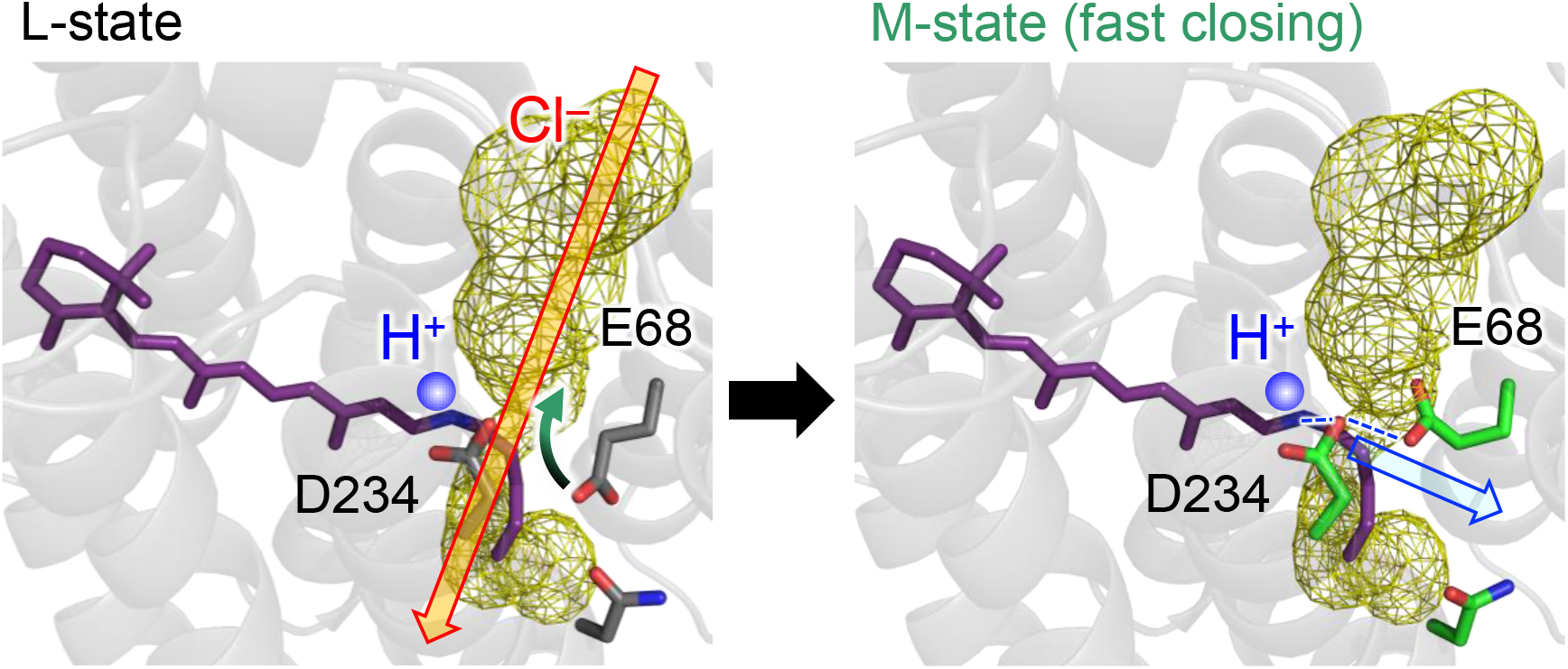
Formation of the proton transfer pathway and disconnection of the anion conduction channel during the L- to M-state transition. The channel is open (red open arrow) in the anion conducting L-state. The reorientation of the Glu68 side-chain (black and green curved arrow) leads to the formation of the H-bond network that proceeds from the Schiff base to Glu68 via Asp234 (blue dotted lines) and the closing of the channel bottle neck (fast channel closing). Thus, the release of the proton from the Schiff base toward Glu68 occurs in the M-state (blue open arrow).

Changes in the protein function, which are caused by mutations of aspartate to asparagine, are often considered as the aspartate being deprotonated in the wild type protein. This can be true if the aspartate is sufficiently far away from the focusing site, where the electrostatic interaction is predominantly approximated by the net charge at the aspartate site. However, this cannot be the case if the aspartate is adjacent to the focusing site (e.g., forming an H-bond), as the H-bond character (e.g., polarity and pattern) of asparagine is not identical to that of protonated aspartate, irrespective of the same net charge. The present example shows that asparagine mutation is not always equivalent to protonated aspartate especially when it is directly involved in the H-bond with the focusing site.

## METHODS

### Coordinates and atomic partial charges

The atomic coordinates were taken from the X-ray structure of *Gt*ACR1 monomer unit “A” (PDB code 6EDQ ^6^). All crystal water molecules were included explicitly in calculations if not otherwise specified. During the optimization of hydrogen atom positions with CHARMM ^24^, the positions of all heavy atoms were fixed, and all titratable groups (e.g., acidic and basic groups) were ionized. The Schiff base was considered protonated. Atomic partial charges of the amino acids and retinal were obtained from the CHARMM22 ^25^ parameter set.

### Protonation pattern

The computation was based on the electrostatic continuum model, solving the linear Poisson-Boltzmann equation with the MEAD program ^26^. The difference in electrostatic energy between the two protonation states, protonated and deprotonated, in a reference model system was calculated using a known experimentally measured p*K*_a_ value (e.g., 4.0 for Asp ^27^). The difference in the p*K*_a_ value of the protein relative to the reference system was added to the known reference p*K*_a_ value. The experimentally measured p*K*_a_ values employed as references were 12.0 for Arg, 4.0 for Asp, 9.5 for Cys, 4.4 for Glu, 10.4 for Lys, 9.6 for Tyr, ^27^, and 7.0 and 6.6 for the N_ε_ and N_δ_ atoms of His, respectively ^28–30^. All other titratable sites were fully equilibrated to the protonation state of the target site during titration. The dielectric constants were set to 4 inside the protein and 80 for water. All water molecules were considered implicitly. All computations were performed at 300 K, pH 7.0, and with an ionic strength of 100 mM. The linear Poisson-Boltzmann equation was solved using a three-step grid-focusing procedure at resolutions of 2.5, 1.0, and 0.3 Å. The ensemble of the protonation patterns was sampled by the Monte Carlo (MC) method with the Karlsberg program ^31^. The MC sampling yielded the probabilities [protonated] and [deprotonated] of the two protonation states of the molecule.

### MD simulations

The *Gt*ACR1 assembly was embedded in a lipid bilayer consisting of 258 1-palmitoyl-2-oleyl-sn-glycero-3-phosphocholine (POPC) molecules using CHARMM-GUI ^32^, and soaked in 29070–29072 TIP3P water models, and 5–7 chloride ions were added to neutralize the system using the VMD plugins ^33^. After structural optimization with position restraints on heavy atoms of the *Gt*ACR1 assembly, the system was heated from 0.1 to 300 K over 5.5 ps with time step of 0.01 fs, equilibrated at 300 K for 1 ns with time step of 0.5 fs, and annealed from 300 to 0 K over 5.5 ps with time step of 0.01 fs. To avoid the influence of changes in the retinal Schiff base structure on the excitation energy, the position restraints on heavy atoms of side-chains were released and MD simulations were performed; the system was heated from 0.1 K to 300 K over 5.5 ps with time step of 0.01 fs and equilibrated at 300 K for 1 ns with time step of 0.5 fs. The system was equilibrated at 300 K for 5 ns with time step of 1.0 fs, and a production run was conducted over 1 ns with 1.0 fs step for sampling of side-chain orientations. All MD simulations were conducted with the CHARMM22 ^25^ force field parameter set using the MD engine NAMD version 2.11 ^34^. For MD simulations with time step of 1.0 fs, the SHAKE algorithm for hydrogen constraints was employed ^34^. For temperature and pressure control, the Langevin thermostat and piston were used ^35,36^. Using the resulting coordinates, the protonation state of the titratable residues was finally determined with the MEAD ^26^ and Karlsberg ^31^ programs.

### QM/MM calculations

After determining the protonation states of all titratable sites and using 10 MD-generated protein conformations, the HOMO-LUMO energy gaps were calculated. The geometry was optimized using a QM/MM approach. The restricted density functional theory (DFT) method was employed with the B3LYP functional and LACVP* basis sets using the QSite ^37^ program. The QM region was defined as the

retinal and Schiff base (Lys238). All atomic coordinates were fully relaxed in the QM region, and the protonation pattern of titratable residues was implemented in the atomic partial charges of the corresponding MM region. In the MM region, the positions of H atoms were optimized using the OPLS2005 force field ^38^, while the positions of the heavy atoms were fixed.

The absorption energy of microbial rhodopsins is highly correlated with the energy difference between highest occupied molecular orbital (HOMO) and lowest unoccupied molecular orbital (LUMO) of the retinal Schiff base (Δ*E*_HOMO-LUMO_) ^4,39^. To calculate absorption energies and corresponding wavelengths, the energy levels of HOMO and LUMO were calculated. The absorption energy (*E*_abs_ in eV) was calculated using the following equation, which is obtained for wild type and six mutant *Gt*ACR1s (coefficient of determination *R*^2^ = 0.93) ^39^:

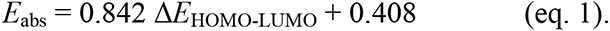

A QM/MM approach utilizing the polarizable continuum model (PCM) method with a dielectric constant of 78 for the bulk region, in which electrostatic and steric effects created by a protein environment were explicitly considered in the presence of bulk water, was employed. Here, the polarizable amber-02 force field ^40^ was applied for the MM region, where the induced dipoles of the MM atoms were considered to reproduce the dielectric screening (i.e., polarizable QM/MM/PCM ^41^). In the PCM method, the polarization points were placed on the spheres with a radius of 2.8 Å from the center of each atom to describe possible water molecules in the cavity. The radii of 2.8–3.0 Å from each atom center and the dielectric constant values of ∼80 are likely to be optimal to reproduce the excitation energetics, as evaluated for the polarizable QM/MM/PCM approach ^41^. The DFT method with the B3LYP functional and 6-31G* basis sets was employed using the GAMESS program ^42^. The HOMO-LUMO energy gap was calculated based on 10 MD-generated protein conformations. See Supporting Information for the QM/MM-optimized atomic coordinates. The electrostatic contribution of the side-chain in the MM region to the absorption wavelength of the retinal Schiff base was obtained from the shift in the HOMO-LUMO energy gap upon the removal of the atomic charges of the focusing side-chain.

### Gene preparation

The cDNA of *Gt*ACR1 (Genbank accession number: KP171708) was optimized for human codon usage and fused to a C-terminal sequence encoding a hexahistidines-tag. The fusion product was inserted into the pCAGGS mammalian expression vector, as previously described ^21,43^. *Gt*ACR1 cDNAs containing mutations were constructed using the In-Fusion Cloning Kit according to the manufacturer’s instructions ^21,43^.

### Protein expression and purification of *Gt*ACR1

The expression plasmids were transfected into HEK293T cells using the calcium-phosphate method ^21,43^. After 1 day incubation, all-*trans*-retinal (final concentration = 5 μM) was added to transfected cells. After another day incubation, the cells were collected by centrifugation (6500 × *g* for 10 min) at 4 °C and suspended in Buffer-A (50 mM HEPES (pH 7.0) and 140 mM NaCl). All-*trans*-retinal (final concentration = 0.31 μM) was added to the cell suspension to reconstitute the photoactive pigments by shaking rotatory for more than 12 hrs at 4 °C. Then, the cells were collected by centrifugation (2900 × *g* for 30 min) at 4 °C and suspended in Buffer-A and solubilized in Buffer-B (20 mM HEPES (pH 7.4), 300 mM NaCl, 5 % glycerol and 1% dodecyl maltoside (DDM)). The solubilized fraction was collected by ultracentrifugation (169800 × *g* for 20 min) at 4 °C and the supernatant was applied to a Ni^2+^ affinity column to purify the pigments. After the column was washed with Buffer-C (20 mM HEPES (pH 7.4), 300 mM NaCl, 5 % glycerol, 0.02% DDM and 20 mM imidazole), the pigment was eluted with a linear gradient of imidazole by Buffer-D (20 mM HEPES (pH 7.4), 300 mM NaCl, 5 % glycerol, 0.02% DDM and 1 M imidazole). Purified samples were concentrated by centrifugation using an Amicon Ultra filter (30000 M_w_ cut-off; Millipore, USA) and the buffer was exchanged using PD-10 column (GE Healthcare, USA) to Buffer-E (20 mM HEPES (pH 7.4), 300 mM NaCl, 5 % glycerol and 0.02 % DDM).

### Spectroscopic analysis

Absorption spectra of the purified proteins were recorded with a UV–visible spectrophotometer (Shimadzu, UV-2450, UV-2600) in Buffer-E. The samples were kept at 15 °C using a thermostat.

## ACKNOWLEDGMENT

This research was supported by AMED (20dm0207060h0004) to Y.S. and JST CREST (JPMJCR1656 to Y.S. and H.I.), JSPS KAKENHI (JP21K15054 to K.K., JP20K21482, JP21H0040413, and JP21H0244613 to Y.S., and JP18H05155, JP18H01937, JP20H03217, and JP20H05090 to H.I.), and the Interdisciplinary Computational Science Program in CCS, University of Tsukuba.

## Competing financial interests

The authors declare no financial and non-financial competing interests.

## Supporting Information

**Figure S1.**
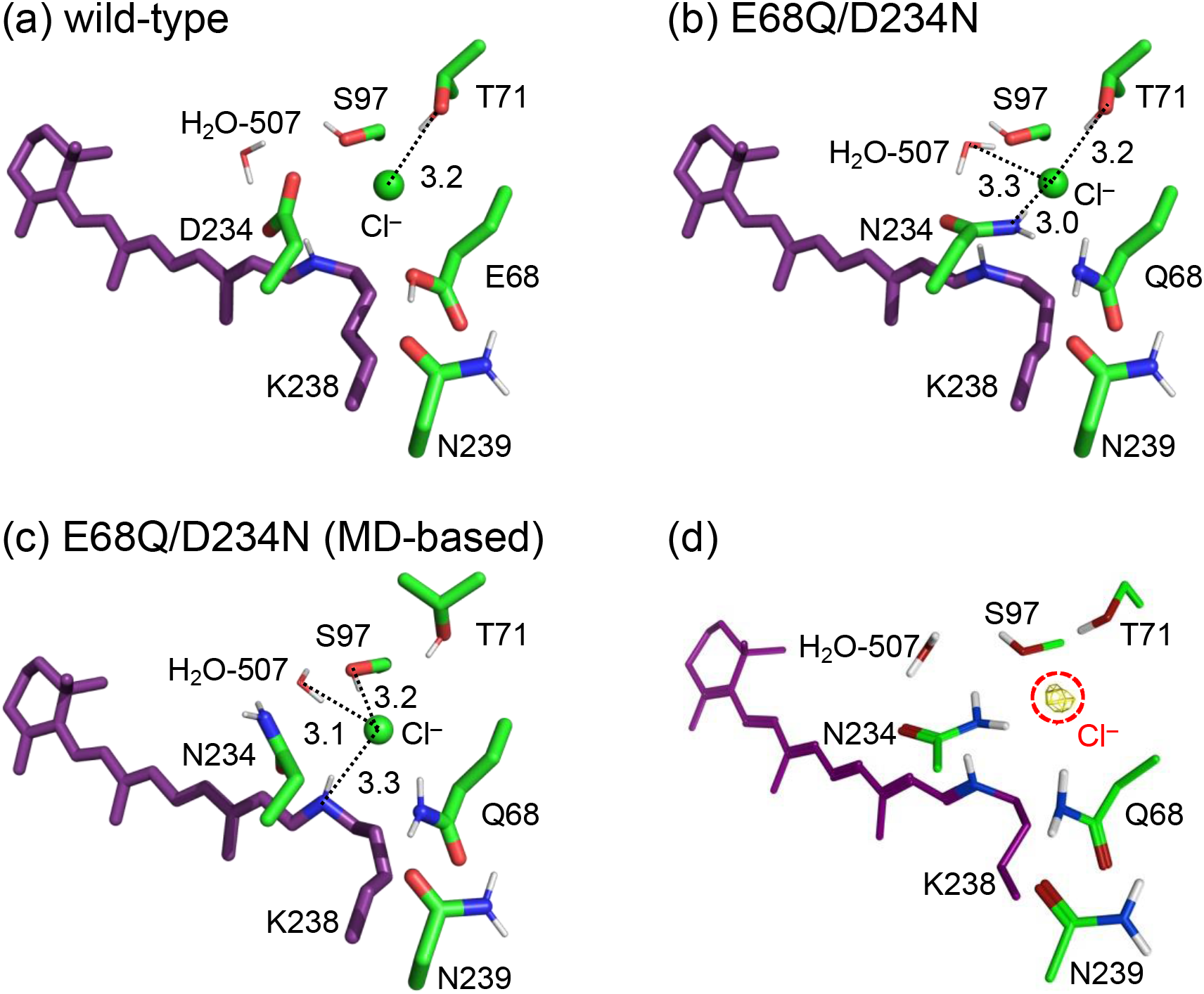
(a, b) QM/MM-optimized Cl^−^-binding structures of (a) wild-type and (b) E68Q/D234N mutant *Gt*ACR1s. For the QM/MM-optimization, the QM region was defined as retinal, the Schiff base (K238), Cl^−^, H_2_O-507, and the side-chains of the residues near the Schiff base (Glu (Gln) 68, Thr71, Tyr72, Ser97, Thr101, Tyr207, Asp (Asn) 234, and Asn239). (c) MD-based Cl^−^-binding E68Q/D234N mutant structure. (d) Distribution pattern of Cl^−^ (yellow mesh) in the QM/MM-optimized E68Q/D234N *Gt*ACR1 structure calculated using a three-dimensional reference interaction site model (3D-RISM). 3D-RISM calculation was performed using the Molecular Operating Environment (MOE) program ^1^.

**Table S1.** Calculated p*K*_a_ values of Glu68 and Asp234 (PDB codes 6EDQ ^2^ and 6CSM ^3^).

|  | **6EDQ** | **6CSM** |
| --- | --- | --- |
| Glu68 | 12 | 9.8 |
| Asp234 | −4.9 | −4.3 |

**Table S2.** Binding energies between Cl^−^ and the surrounding environments in Cl^−^-binding wild-type and E68Q/D234N mutant structures (kcal/mol). The binding energies were calculated based on the QM/MM-optimized and MD-based structures, using the MOE program ^1^.

|  | wild-type | E68Q/D234N | E68/D234N<br>(MD-based) |
| --- | --- | --- | --- |
| Thr71 | -3.9 | -3.8 | – |
| Ser97 | – | – | -4.1 |
| Asn234 | – | -7.7 | – |
| Lys238<br>(Schiff base) | -2.9 | -1.9 | -12.7 |
| H<sub>2</sub>O-507 | – | -3.0 | -3.5 |
| Total | -6.8 | -16.4 | -20.3 |

## Notes

### Competing Interest Statement

The authors have declared no competing interest.

